# AtlasLens: Metadata-centric exploration and analysis of single-cell atlases

**DOI:** 10.64898/2026.07.10.735823

**Authors:** Shamim Ashrafiyan, Iaroslav Kosaretskii, Marcel H Schulz

## Abstract

**Motivation:** The rapid expansion of single-cell RNA sequencing (scRNA-seq) atlases has generated datasets comprising millions of cells annotated with increasingly rich metadata, including tissue, cell type, disease status, sex, age, treatment, and temporal information. Biological questions frequently require simultaneous interrogation of multiple metadata dimensions, such as identifying specific cell populations within defined tissues, disease states, demographic groups, and time points. While existing interactive platforms facilitate visualization and analysis of scRNA-seq data, deep metadata-driven exploration and downstream analysis of atlas-scale datasets remain insufficiently supported.

**Results:** We developed AtlasLens, an open-source R/Shiny application for interactive exploration of scRNA-seq datasets and integrated cellular atlases. AtlasLens enables iterative filtering across arbitrary metadata combinations, allowing users to define biologically meaningful cellular subsets and immediately perform downstream analyses. The platform integrates interactive visualization, differential expression analysis, Gene Ontology enrichment with redundancy reduction, temporal expression analysis, and context-dependent gene function profiling through GeneCOCOA. AtlasLens additionally records analysis history and automatically generates corresponding R code to enhance reproducibility. The application is distributed through Docker for simple local deployment, preserving data privacy and eliminating dependency-management challenges. We demonstrate AtlasLens using the Tabula Muris and a time-resolved whole-lung single-cell atlas of bleomycin-induced lung injury and fibrosis, highlighting its ability to support complex metadata-driven biological investigations.

**Availability:** Source code is available at https://github.com/SchulzLab/AtlasLens.

## Introduction

Single-cell RNA sequencing (scRNA-seq) has enabled high-resolution characterization of cellular heterogeneity across tissues, developmental stages, and disease states, leading to the generation of increasingly large cellular atlases. Major initiatives such as the Human Cell Atlas (HCA)(Regev et al., 2017), Tabula Sapiens (Tabula Sapiens Consortium, 2022), Tabula Muris (Tabula Muris Consortium, 2018), and numerous organ-specific atlases now provide rich, large-scale datasets containing complex cellular, temporal, and phenotypic metadata. Crucially, these resources are not merely large in cell number; they encode multiple intersecting biological dimensions, including cell type, tissue of origin, disease status, sex, age, and time point. As scRNA-seq atlases continue to grow, biological discovery increasingly depends on the ability to interrogate specific cellular subsets defined by combinations of metadata attributes rather than by individual variables alone. For example, a researcher studying cardiac remodeling may wish to investigate endothelial cells from aged female animals at a particular time point following myocardial infarction, compare them to matched controls, and subsequently perform differential expression, pathway enrichment, or gene functional analyses. Consequently, there is a growing need for tools that support flexible, multi-dimensional metadata-driven interrogation of single-cell atlases together with downstream analytical interpretation.

While existing platforms have substantially improved accessibility of single-cell datasets, current solutions generally fall into two categories. The first category comprises visualization and data exploration tools: dataset browsers and framework-based viewers focused on interactive inspection of single-cell data. Atlas browsers such as cellxgene (Megill et al., 2021), as well as framework-based viewers such as ShinyCell (Ouyang et al., 2021) and iSEE (Rue-Albrecht et al., 2018) enable efficient exploration of cellular embeddings, gene expression patterns, and metadata annotations, but offer limited support for downstream analyses. The second category comprises analytical platforms that provide functional analysis capabilities. For example, ASAP (Gardeux et al., 2017) offers analytical functions, while SCHNAPPs (Jagla et al., 2021), ICARUS v3 (Jiang et al., 2024), and ScRDAVis (Jagadesan and Guda, 2025) additionally perform extensive preprocessing and analysis. These tools, however, are often focused on complete analysis pipelines rather than iterative and deep filtration of existing atlases. Consequently, most existing platforms provide limited support for simultaneous combinatorial filtering across multiple metadata dimensions, forcing researchers to define cellular subsets using sequential or restricted filtering strategies, which hinders complex cohort definition and downstream analysis (Table S1 compares AtlasLens with existing tools).

This limitation reflects a fundamental gap in current single-cell analysis ecosystems: the lack of support for flexible combinatorial querying across multiple metadata dimensions, coupled with integrated visualization and downstream analytical functionality within a unified environment. To address this challenge, we developed AtlasLens, a containerized R/Shiny application specifically designed for metadata-centric exploration of integrated scRNA-seq atlases. AtlasLens bridges the two categories, combining interactive visualization and data exploration with integrated downstream analysis. In detail, it supports flexible integration of user-defined metadata and enables iterative filtering across arbitrary combinations of biological and experimental metadata variables, allowing researchers to define biologically meaningful cellular subsets for downstream analyses. AtlasLens seamlessly couples subset selection with differential expression analysis, Gene Ontology enrichment with redundancy reduction, temporal gene-expression analysis, and context-dependent gene function profiling. By combining atlas-scale metadata interrogation, integrated biological interpretation, local deployment, and reproducibility support within a unified framework, AtlasLens facilitates efficient exploration of complex single-cell atlas resources.

### Implementation

AtlasLens is built entirely in R using the Shiny framework, leveraging Seurat objects as the data structure, with two scripts provided in the repository to convert AnnData or related files into Seurat. The Seurat object is supplied at startup via an environment variable, and AtlasLens automatically detects the relevant metadata columns; an optional JSON configuration file can override these assignments, set the species, and provide personalized descriptive content for the landing page. For accessibility and reproducibility, the full environment is distributed as a Docker container. Users can download or clone the GitHub repository, build the container, and launch the app locally without dependency conflicts or data transfer to remote servers.

### Features

The key feature of the AtlasLens is deep multi-dimensional metadata filtering, allowing users to iteratively subset cells using combinations of metadata variables to answer specific biological questions and investigate the cellular subpopulations. This feature is directly supported in all analysis and visualization panels through the app.

#### Interactive data exploration and visualization

AtlasLens provides interactive exploration of scRNA-seq atlases and datasets through dynamic UMAP visualization and metadata-driven filtering. Cells can be colored according to any available metadata attribute. Side-by-side UMAP visualization enables comparison between selected metadata distributions and gene expression patterns, as well as co-expression analysis of two selected genes. Gene expression patterns of a group of selected genes can additionally be explored using dot plots across selected metadata groups.

Moreover, AtlasLens automatically fetches total transcript counts (nCount), detected features (nFeature) and calculates the mitochondrial transcript percentage (percent.mt) for the filtered metadata and allows users to visualize them to assess and check the quality of the data.

#### Differential expression and enrichment

The app supports metadata-based differential expression analysis with interactive result tables and volcano plot visualization. Users can define custom thresholds for adjusted p-values and log fold-change values, and selected genes can be highlighted directly on volcano plots for rapid inspection. Gene Ontology enrichment analysis (Yu et al., 2012) is integrated to facilitate biological interpretation of differentially expressed genes. As an important feature for summarizing enrichment results, the app incorporates the R package rrvgo (Sayols, 2023) to reduce redundancy among overlapping GO terms. Similar GO terms are clustered based on semantic similarity and visualized using scatter plots and treemaps, providing a clearer and more interpretable overview of enriched biological processes.

#### Time-series analysis

For datasets with temporal structure, the application provides advanced visualization and clustering of gene expression dynamics across time points. Users can visualize gene expression using boxplots and violin plots across time points, identify temporal expression patterns using k-means clustering, and generate heatmaps summarizing average expression per time point. Specifically, genes are grouped into clusters based on their expression profiles across time points, enabling the identification of genes with similar temporal dynamics. This allows users to detect coordinated biological programs, such as early-response or late-response gene modules. This combination of visualization and clustering provides both descriptive and pattern-based insights, which are not commonly integrated in existing interactive scRNA-seq tools.

#### Gene function analysis

AtlasLens integrates GeneCOCOA-based context-dependent gene function analysis (Zehr et al., 2025) directly into an interactive exploration workflow. Unlike conventional gene set enrichment analyses that begin with a gene set, GeneCOCOA starts from a single gene and uses dataset-specific co-expression patterns to infer the biological pathways and functions most strongly associated with that gene. Users can select a gene of interest, define biological conditions through metadata-based filtering (e.g., healthy versus disease states), and compare associated functional pathways across selected groups. Results are visualized as comparative bar plots, enabling identification of condition-specific functional and co-regulatory patterns.

#### Reproducibility and history tracking

To improve reproducibility, AtlasLens maintains a session-based analysis history that allows users to revisit previous analyses during an active session. For each analysis step, automatically generated R code can be exported, enabling reproduction of analytical workflows. Moreover, the open-source design of the app enables advanced users to extend and customize the application.

## Results

To demonstrate the capabilities of AtlasLens, we used the publicly available Tabula Muris dataset, which includes 23 tissues, 21,025 genes, and 110,824 cells (Tabula Muris Consortium, 2018) and whole-lung single-cell RNA-seq time course of bleomycin-induced lung injury (Strunz et al., 2020), consisting of 23,400 genes and 29,297 cells sampled at six timepoints after bleomycin instillation which captures the progression of lung injury, alveolar regeneration, and fibrosis. An overview of the AtlasLens core functionalities is presented in Figure 1a. Figure 1b illustrates metadata-driven UMAP exploration using the Tabula Muris dataset for male mice, with cells colored by tissue of origin across 22 tissues, demonstrating how AtlasLens supports rapid visual orientation within large multi-tissue atlases. Figure 1c shows a summarized GO enrichment visualization produced by rrvgo (Sayols, 2023), where semantically redundant GO terms are clustered and reduced to representative biological themes. Here, enrichment results from a DEA between regular ventricular cardiac myocyte and regular atrial cardiac myocyte for heart tissue in the Tabula Muris dataset are condensed into interpretable process categories such as ‘Heart process’, ‘Cardiac muscle cell potential’, and ‘Mitochondrial RNA metabolic process’, illustrating the value of redundancy reduction for biological interpretation. Figure 1d presents temporal k-means clustering of gene expression dynamics in Krt8+ alveolar differentiation intermediate (ADI) cells across lung injury time points. Genes are grouped into eight clusters based on their expression trajectories, enabling identification of early-response, mid-phase, and late-remodeling gene modules. Users can extract genes from individual clusters for downstream differential expression or GO enrichment analysis. Figure 1e shows cell-type-resolved functional profiling of Krt8 in Krt8 ADI cells across the injury time course (days 3–28 after bleomycin instillation) using GeneCOCOA (Zehr et al., 2025). Krt8 marks the transitional alveolar stem-cell state that emerges during regeneration and persists in lung fibrosis (Strunz et al., 2020). The functions most associated with Krt8 vary substantially across time points, recapitulating the p53 and NF-κB activation and cellular-stress signature of this state, alongside inflammatory and metabolic programs, and reflecting its dynamic reprogramming during lung injury and fibrosis. The detailed plots and functions are shown in supplementary fig S1-S11.

**Figure 1.**
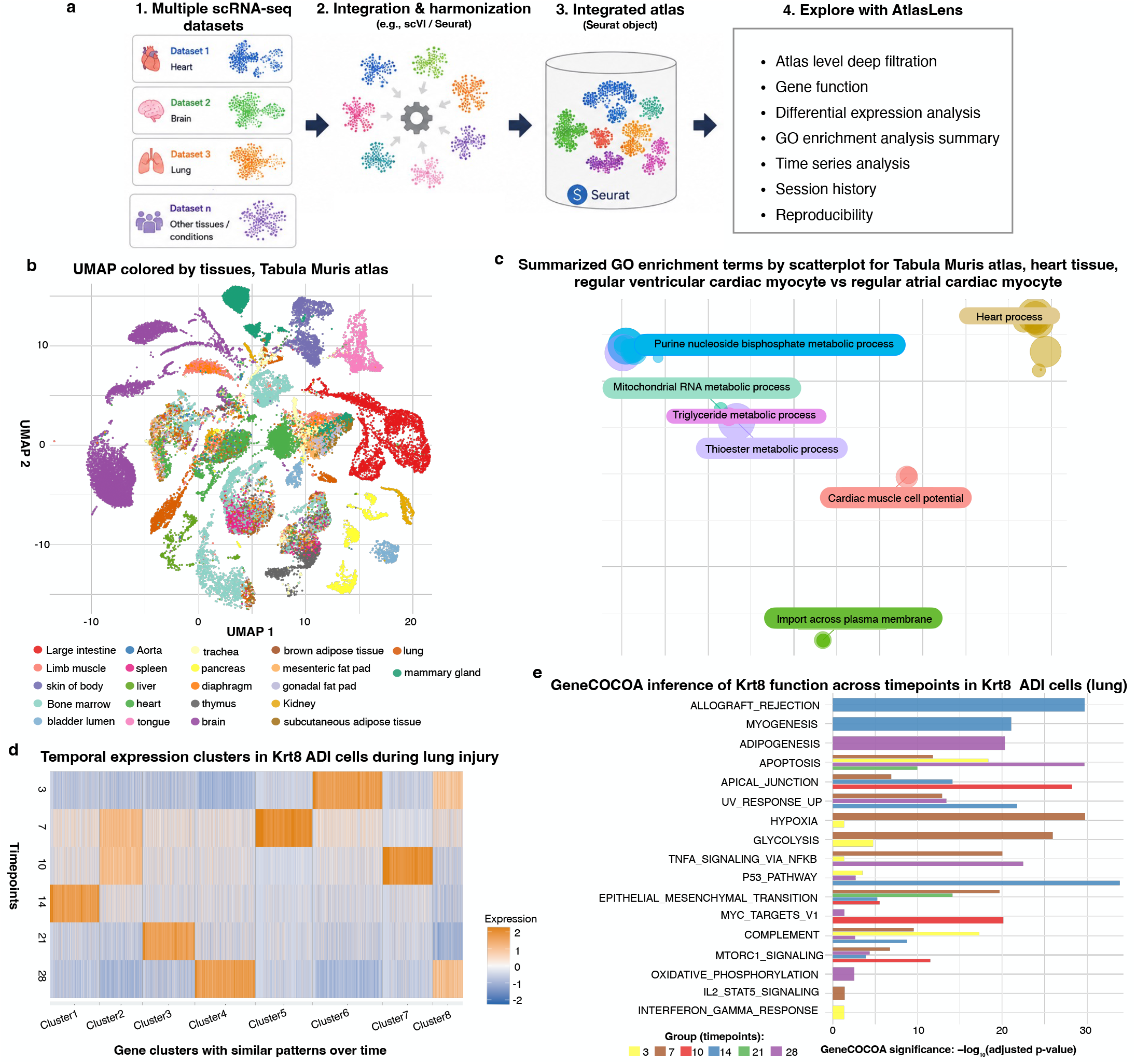
AtlasLens workflow overview: **(a)** AtlasLens works with both single-cell dataset and atlases containing rich, diverse metadata built using integration tools such as Seurat and scVI (Lopez et al., 2018). To use AtlasLens and its functions (step 4), integrated atlases should be saved as a Seurat object; conversion scripts for AnnData and related formats are provided in the repository. **(b)** Metadata driven UMAP exploration. **(c)** Summarized GO analysis terms shown by scatterplot (more plots are shown in Fig. S7 and S8). **(d)** Temporal clustering of gene expression patterns. **(e)** Functional profiling of a gene using GeneCOCOA.

## Conclusion

AtlasLens addresses a growing need in single-cell genomics for interactive tools capable of exploring atlas-scale datasets with complex, multi-dimensional metadata structures. Within a unified framework, it enables biologists to answer their own questions by filtering metadata at arbitrary levels and immediately performing downstream analysis. It combines capabilities that are often distributed across multiple existing tools, including multidimensional metadata filtering, temporal expression analysis, GO redundancy reduction, context-dependent gene functional profiling, and reproducible workflow tracking. The local deployment model preserves data privacy, while session history and automatic R code generation ensure that analyses remain transparent and reproducible. As single-cell atlases grow in biological scope and metadata complexity, AtlasLens provides a way to handle this complexity and extract meaningful biological insights.

## Author contributions

S.A.: Conceptualization, Software, and Writing original draft. I.K.: Software. M.H.S.: Supervision, project administration, and writing review and editing.

## Supplementary material

Supplementary Table 1 provides a detailed feature comparison of AtlasLens with existing interactive scRNA-seq visualization tools. Supplementary Figures S1–S11 illustrate the full range of AtlasLens functionalities demonstrated in this study.

## Data Availability

Publicly available single-cell RNA sequencing data from the Tabula Muris (Tabula Muris Consortium, 2018) and a whole-lung bleomycin injury time-course dataset (Strunz et al., 2020) were utilized in this study. The processed Seurat objects generated from both datasets have been deposited in Zenodo (Ashrafiyan et al., 2026b). The AtlasLens source code is available on GitHub at https://github.com/SchulzLab/AtlasLens and archived on Zenodo (Ashrafiyan et al., 2026a).

