## Supplementary file for "AtlasLens: Metadata-centric exploration and analysis of single-cell atlases"

**Table 1**: Comparison of different scRNA-seq visualization and analytic tools in terms of their input files, analysis features, and user interface. The exploration and analysis features are described in detail in Section S1.

| Functionality | AtlasLens | ICARUS v3 | SCHNAPPs | ScRDAVis | ASAP | iSEE | ShinyCell | Cellxgene |
|---|---|---|---|---|---|---|---|---|
| **Input file format** | | | | | | | | |
| Seurat object | ✓ | ✓ | ✓ | ✓ | | | ✓ | ✓ |
| H5 files | ✓ | ✓ | | ✓ | | | ✓ | |
| Single matrix file | ✓ | ✓ | ✓ | ✓ | ✓ | | | |
| SummarizedExperiment object | | | | | ✓ | ✓ | | |
| **Preprocessing and annotation** | | | | | | | | |
| Preprocessing (QC filtering, PCA, cell type prediction, Doublet identification, cell-cell communication, cell type annotation...) | | ✓* | ✓* | ✓ | | | | |
| **Exploration and analysis features** | | | | | | | | |
| Marker identification/DEA | ✓ | ✓ | ✓ | ✓ | ✓ | ✓ | ✓ | ✓ |
| QC visualization in selected metadata | ✓ | ✓ | ✓ | ✓ | ✓ | | | |
| Markers between specific clusters | ✓ | | | | | ✓ | | ✓ |
| Showing gene expression (UMAP, heat map or dot plot) | ✓ | ✓ | ✓ | ✓ | ✓ | ✓ | ✓ | ✓ |
| Gene coexpression plot | ✓ | | ✓ | ✓ | | | ✓ | |
| GO enrichment / Pathway / GSEA analysis | ✓ | ✓ | | ✓ | ✓ | | | ✓ |
| Summarized GO visualization | ✓ | | | | | | | |
| Temporal gene expression analysis | ✓ | | | | | | | |
| Function of a gene of interest | ✓ | | | | | | | |
| Showing metadata of each single cell | ✓ | | | | | ✓ | | ✓ |
| Combinatorial metadata filtering at the same time | ✓ | | | | | | | ✓ |
| Session history | ✓ | ✓ | ✓ | | | | | |
| **File export and user interface** | | | | | | | | |
| Reproducibility (generating a script) | ✓ | | ✓ | | | ✓ | | |

| | | | | | | | | |
|---|---|---|---|---|---|---|---|---|
| Working local (no need to upload datasets) | ✓ | ✓ | ✓ | ✓ | | ✓ | ✓ | |
| Saving the results (image & csv) | ✓ | ✓ | ✓ | ✓ | ✓ | ✓ | ✓ | ✓ |
| Command line/GUI | ✓ | ✓ | ✓ | ✓ | | ✓ | ✓ | |
| Language/platform | R | R | R | R | multi | R | R | Python |

✓* Partially supported

AtlasLens is designed specifically as an atlas-level exploration and analysis tool. It accepts pre-processed and pre-integrated Seurat objects as input and does not perform upstream steps such as quality control, doublet removal, or cell-type annotation, since these are handled by dedicated tools; e.g., Seurat (Butler et al., 2018; Stuart et al., 2019) and Scater (McCarthy et al., 2017) for quality control and preprocessing, scDblFinder (Germain et al., 2021) or DoubletFinder (McGinnis et al., 2019) for doublet detection, and SingleR (Aran et al., 2019) or Azimuth (Hao et al., 2021) for cell-type annotation. AtlasLens focuses on flexible, multi-dimensional metadata-driven exploration and integrated biological interpretation of atlas-scale datasets.

### S1. Features in the exploration and analysis category

**QC visualization in selected metadata:** The tools provide QC metrics and visualizations such as number of detected genes (nFeature), total counts per cell (nCount), and percentage of mitochondrial genes. AtlasLens shows three QC for any level of filtration, users can see the plots for different metadata, for example they can plot them for specific cell types or different dataset or tissue and even a group of filtered metadata (Fig. S1a)

**Marker identification / DEA and Markers between specific clusters:** Identifies marker genes and performs Differential Expression Analysis (DEA) using wilcoxon test between clusters, cell types, or experimental conditions. AtlasLens does DEA for any combination of metadata and deep filtration, for example users can choose sub datasets and dataset specific cell type, sex and more metadata for this analysis. In this section we show a volcano plot and table of the results and users are able to set the p-value and log fold change values and download the results and the plot (Fig. S2)

**UMAP / t-SNE visualization:** Provides low-dimensional visualization of single-cell data using UMAP or t-SNE. AtlasLens shows two UMAP plots next to each other which one shows the selected metadata and the other one shows expression of a selected gene in the selected metadata (Fig. S3)

**Showing gene expression (heat map/dot plot/violin plot):** Visualizes expression levels of selected genes across clusters or cell types using heatmaps, dot plots, violin plots. In addition to UMAP visualization, AtlasLens provides a dot plot feature for user-selected or uploaded gene lists. This allows users to explore gene expression across chosen metadata categories, such as cell type or gender, enabling flexible comparison of expression patterns across different biological groups (Fig. S4)

**Gene coexpression plots:** Showing expression of two genes at the same time on the low dimensional embeddings is useful and helps to understand their pattern on a specific metadata like cell type (Fig. S5).

**Gene Ontology enrichment / Pathway / GSEA analysis:** Performs functional enrichment analyses including Gene Ontology (GO), pathway enrichment, or Gene Set Enrichment Analysis (GSEA) to interpret biological functions and pathways associated with gene sets. AtlasLens allows users to select genes from DEA results or upload gene lists, and visualize enrichment results as dot plots (Fig. S6)

**Summarized GO visualization:** In addition to normal visualization of GO terms, AtlasLens summarizes the results using the rrvgo package (Sayols, 2023) and provides two additional summarized views: scatterplot and treemap. Fig. 1c shows the scatterplot for the summarized GO for regular ventricular cardiac myocyte vs regular atrial cardiac myocyte in heart tissue. Fig. S7 shows also the scatterplot for neuron cells vs oligodendrocyte in the brain tissue. Fig. S8 shows the treemap version for the same combination.

**Temporal gene expression analysis:** AtlasLens uniquely enables users to select a dataset from their atlas, specify a cell type of interest, and examine gene expression dynamics across defined time points for individual genes. In addition, it applies k-means clustering to group genes based on their temporal expression profiles, visualized as a heatmap. Users can extract genes from specific clusters for downstream analyses, including DEA and GO enrichment (Fig. 1d and Fig. S9 ).

**Gene function:** AtlasLens enables users to select a gene of interest along with specific metadata conditions (such as tissue type, cell type, and experimental state) to investigate gene function in a context-dependent manner across the rest of the dataset. For example, within a selected tissue like lung, a selected cell type such as Krt8 ADI (alveolar differentiation intermediate) cells, and time points such as (3, 7, 10, 14, 21, and 28), AtlasLens allows users to explore how the functional role and associations of a gene (like Krt8) vary across multiple time points, which may extend beyond a simple binary comparison. This analysis is implemented using the GeneCOCOA package (Zehr et al., 2025), which supports context-aware functional characterization of genes based on co-expression patterns (Fig. 1e)

**Showing metadata of each single cell:** AtlasLens provides detailed metadata information for individual single cells. Users can select specific cells directly from the UMAP embeddings and inspect all associated metadata for the selected cell. To facilitate precise cell selection, AtlasLens also supports interactive zooming and navigation within UMAP visualizations (Fig. S10)

**Advanced metadata filtration (Atlas level):** Enables advanced and multi-level filtering of cells simultaneously based on complex metadata combinations across rich metadata atlases, facilitating precise subset selection and exploration.

**Session history:** AtlasLens provides a dedicated *History* panel that keeps track of analyses generated during an active session. Users can interactively revisit previous results, reopen plots and analyses, and download outputs for further use and reproducibility (Fig. S11).

### Figures:

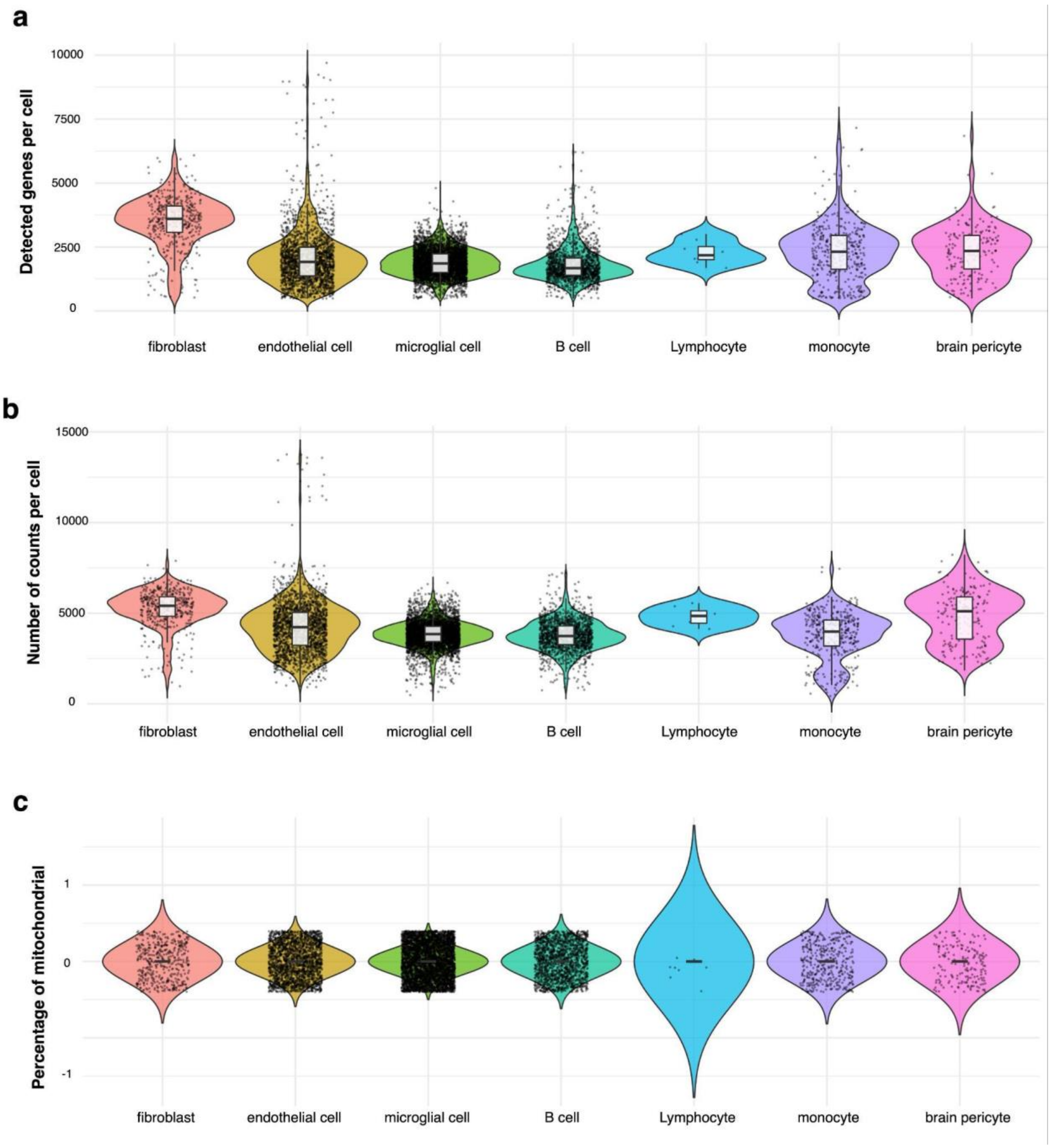


**Fig. S1**. **Quality control metrics for female cells across selected cell types in the Tabula Muris dataset.** (a) Number of detected features per cell (nFeature). (b) Total transcript counts per cell (nCount). (c) Percentage of mitochondrial gene expression (percent.mt). AtlasLens enables QC visualization at any level of metadata filtration, allowing users to inspect metric distributions within specific subpopulations.

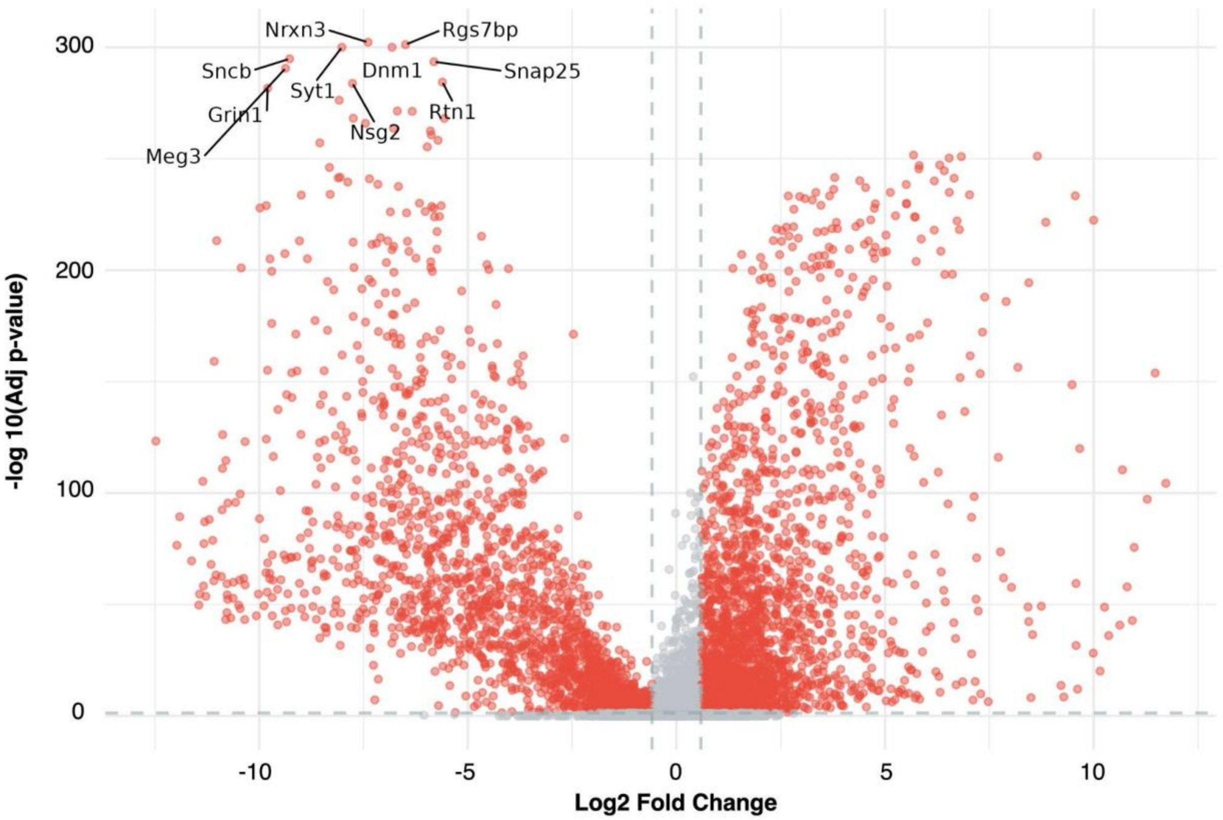


**Fig. S2. Differential expression analysis between neuron and oligodendrocyte cells in brain tissue from the Tabula Muris dataset.** The volcano plot displays log fold-change against adjusted p-values; users can interactively adjust significance thresholds and highlight genes of interest. The list of significant differentially expressed genes can be downloaded. The differential expression tab produces a table with a full list of genes, p-values and log-fold change values which is also downloadable.

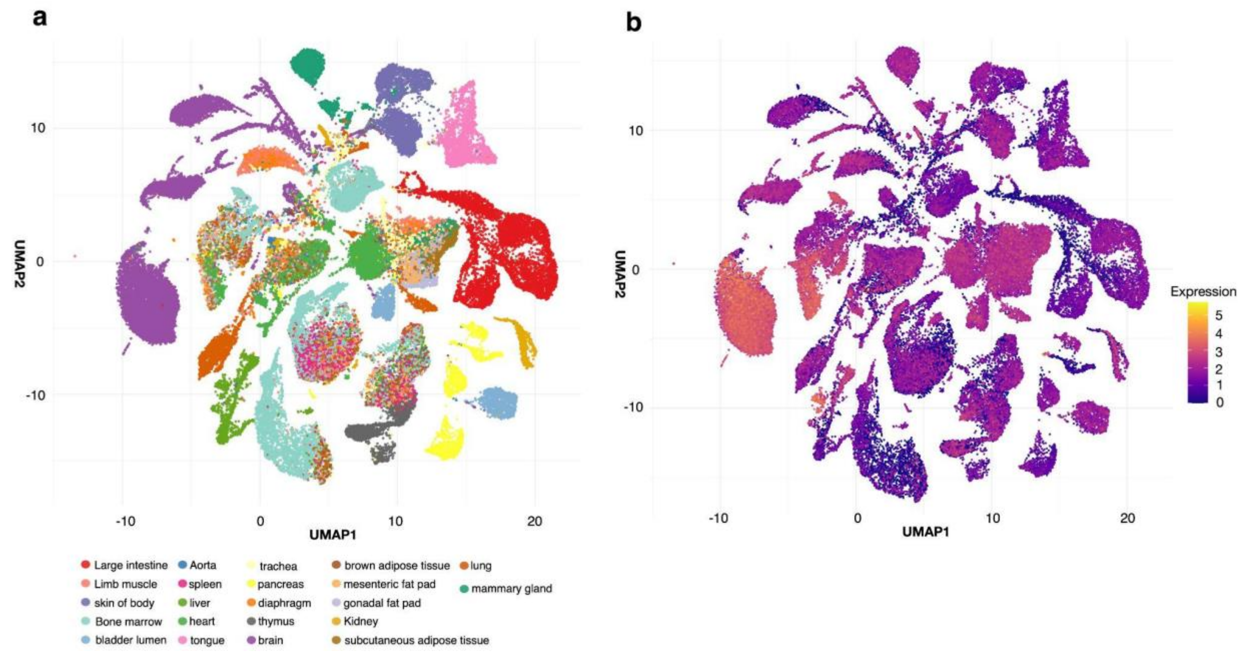


**Fig. S3. Side-by-side UMAP visualization for metadata and gene expression exploration in the Tabula Muris dataset.** (a) UMAP of cells from mice aged 3 and 18 months, colored by tissue of origin. (b) Expression of the gene *Tpm2* (ENSMUSG00000021939) projected onto the same embedding, enabling direct comparison of expression patterns with metadata distributions. As shown on the right plot, *Tpm2* shows higher expression in brain tissue in this dataset.

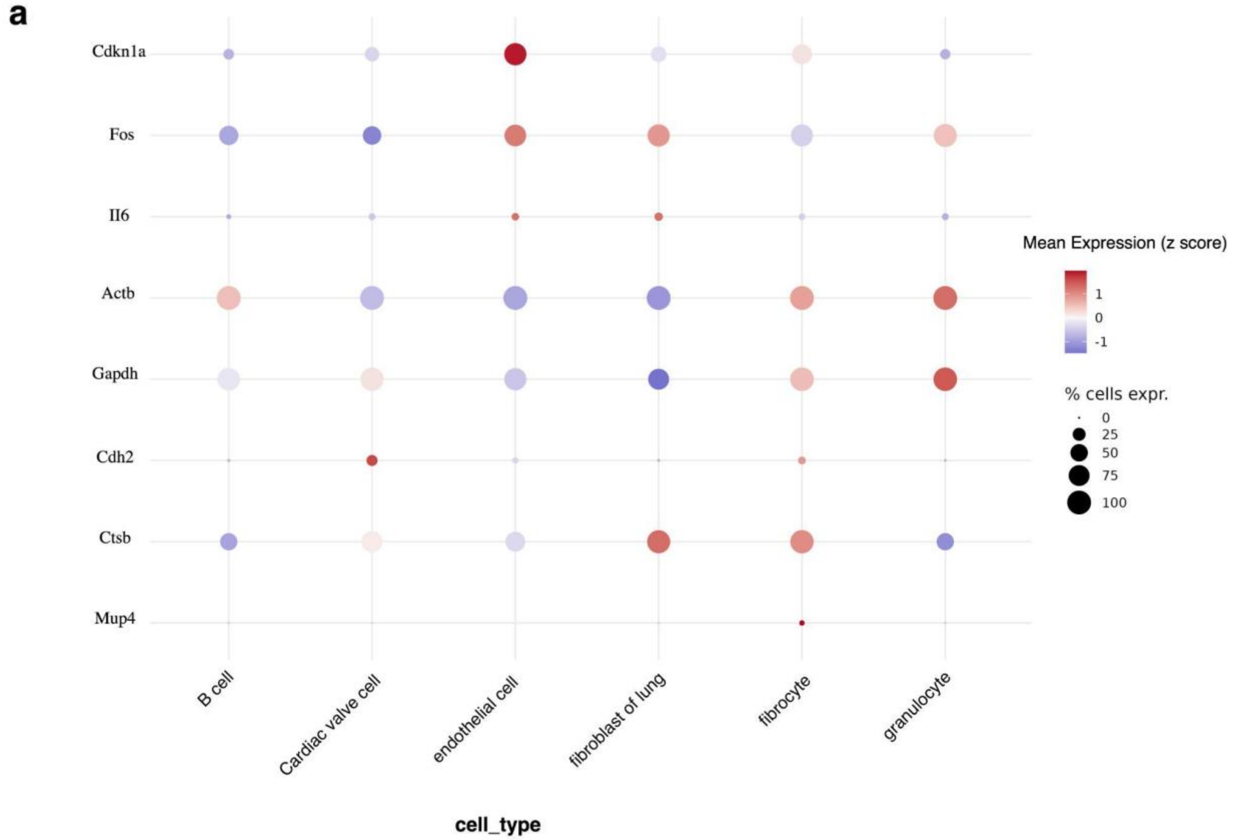


**Fig. S4. Dot plot of gene expression across selected cell types (Tabula Muris, male mice).** Dot color shows mean log-normalized expression z-scored across the displayed cell types (per gene); dot size shows the percentage of cells expressing each gene. Genes and metadata groupings are flexible and user-defined.

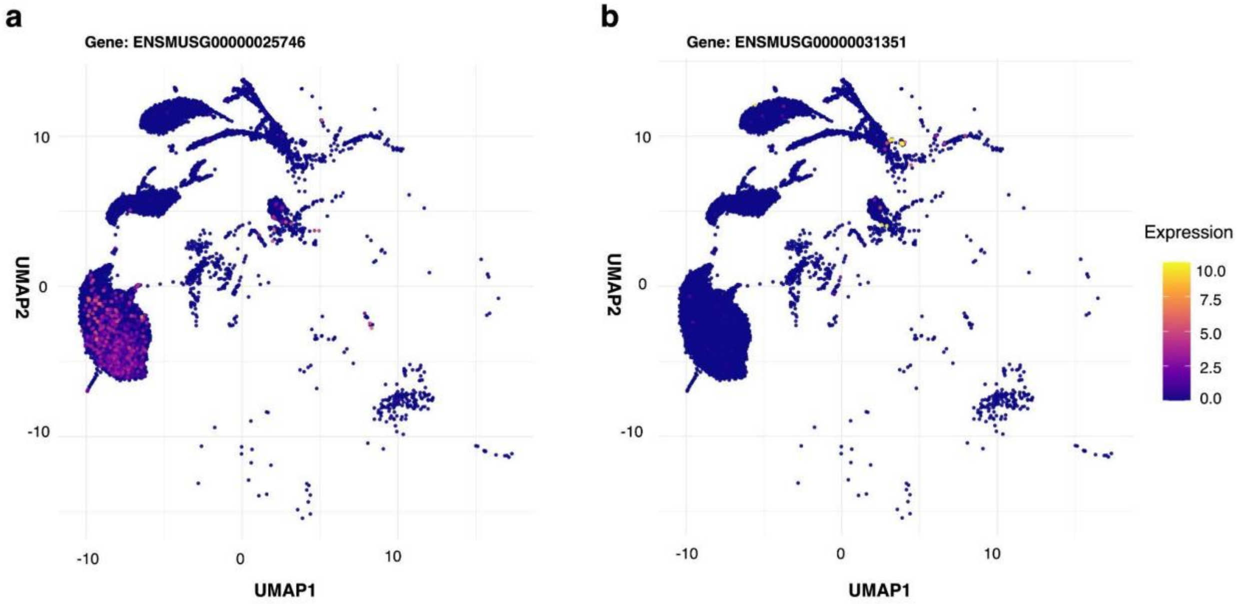

**Fig. S5. Gene co-expression visualization in brain tissue from the Tabula Muris dataset.** (a) Expression of *Cntnap2* (ENSMUSG00000025746) which has higher expression than the *Nrxn1* (ENSMUSG00000031351) gene shown on panel (b). Side-by-side display of two genes on the same UMAP embedding supports visual assessment of spatial co-expression patterns within a defined metadata context.

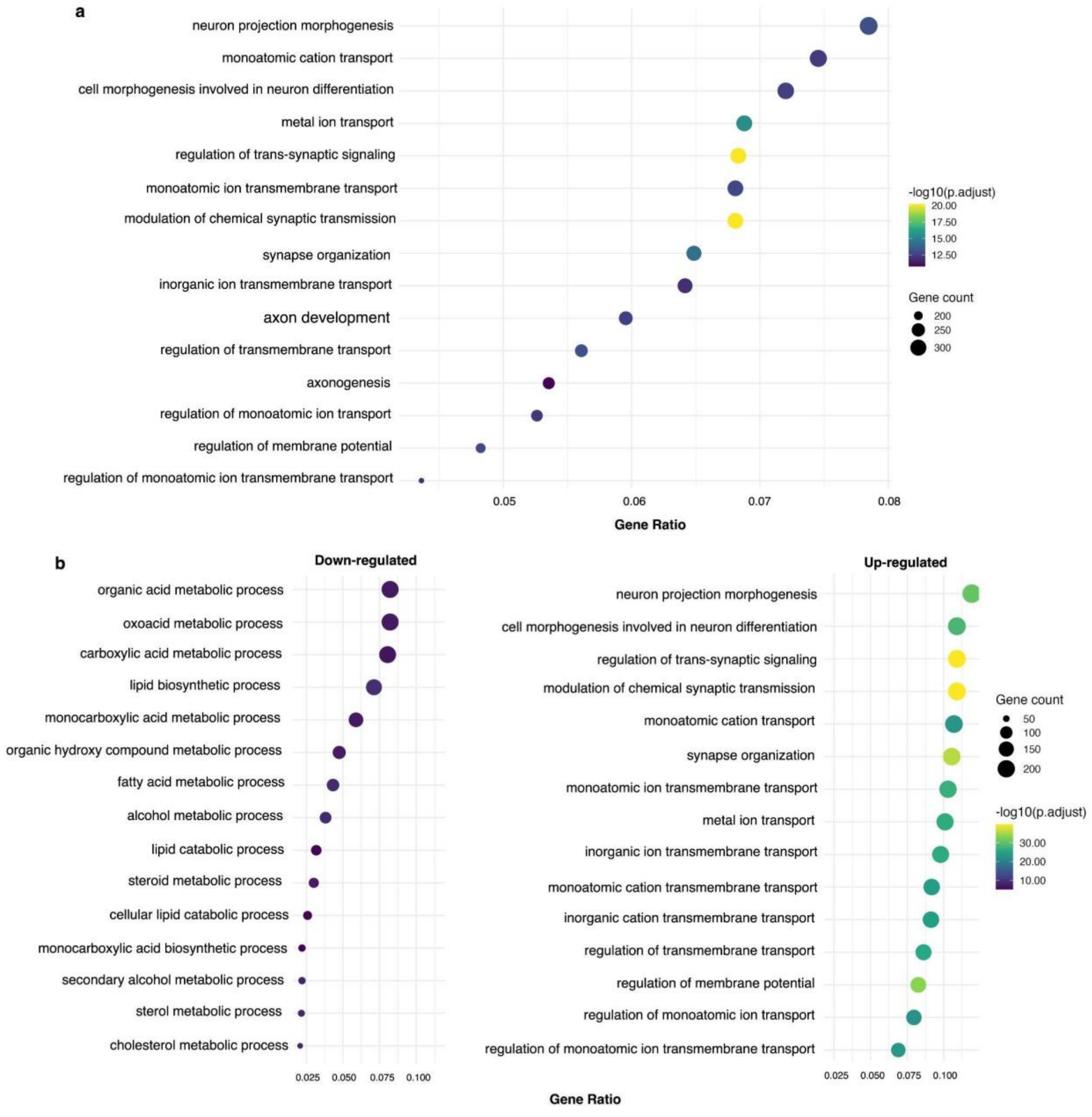


**Fig. S6. Gene Ontology (GO) enrichment analysis of genes differentially expressed between oligodendrocytes and neurons in mouse brain tissue (Tabula Muris).** Differential expression was computed with neurons as the reference, so positive log2 fold-changes denote higher expression in oligodendrocytes. **(a)** GO enrichment for all significant differentially expressed genes (DEGs). **(b)** GO enrichment stratified by direction of regulation: genes upregulated in oligodendrocytes (right) and downregulated in oligodendrocytes, i.e. enriched in neurons (left). In all panels, dot size indicates gene ratio and color reflects the adjusted *p*-value. Genes can be drawn from the differential expression results or supplied as a custom list.

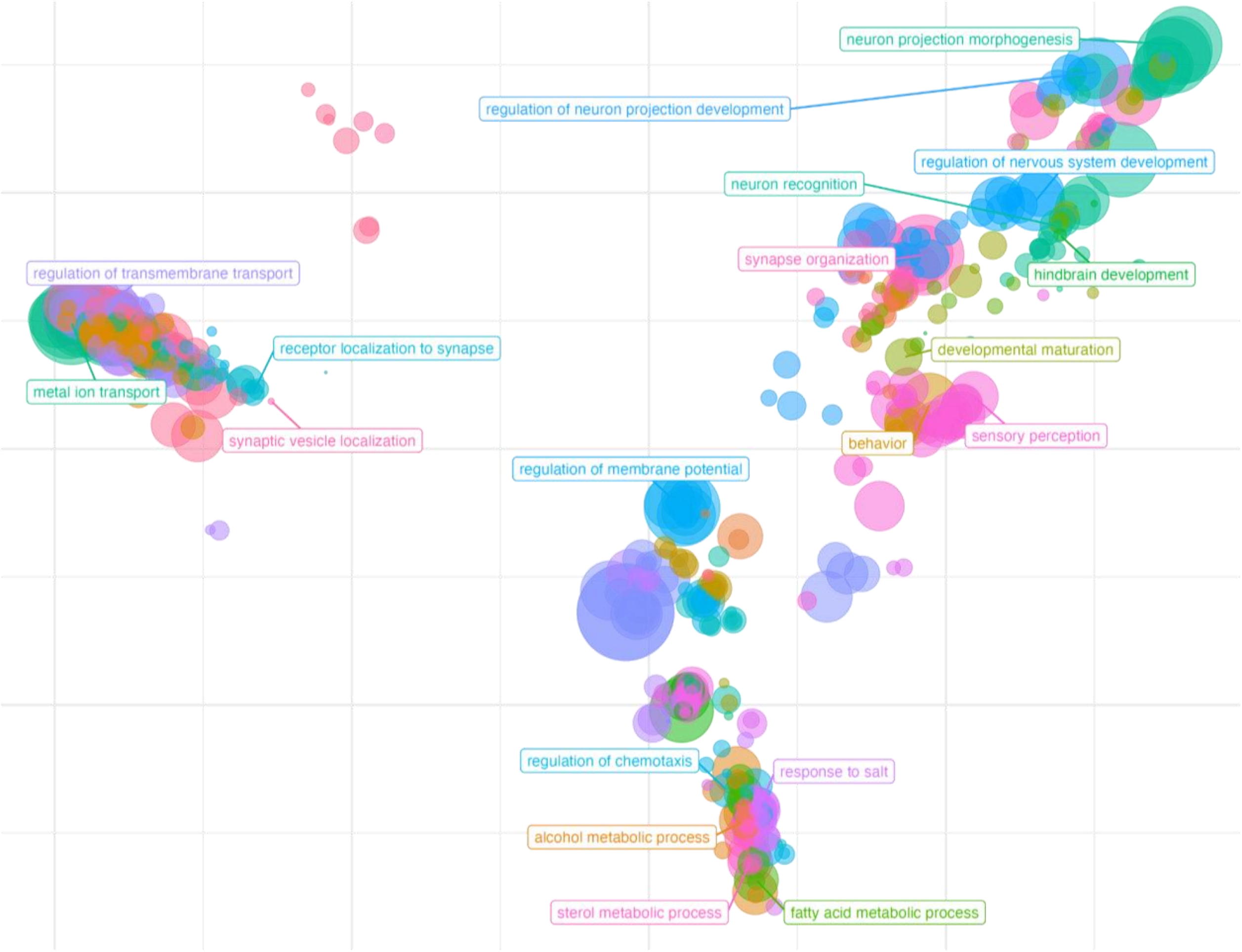


**Fig. S7. GO enrichment analysis summarization by scatterplot:** scatterplot representation of pairwise semantic similarity between enriched GO terms, corresponding to the enrichment results for oligodendrocytes cells versus neurons cells in brain tissue which is shown in Fig. S6.

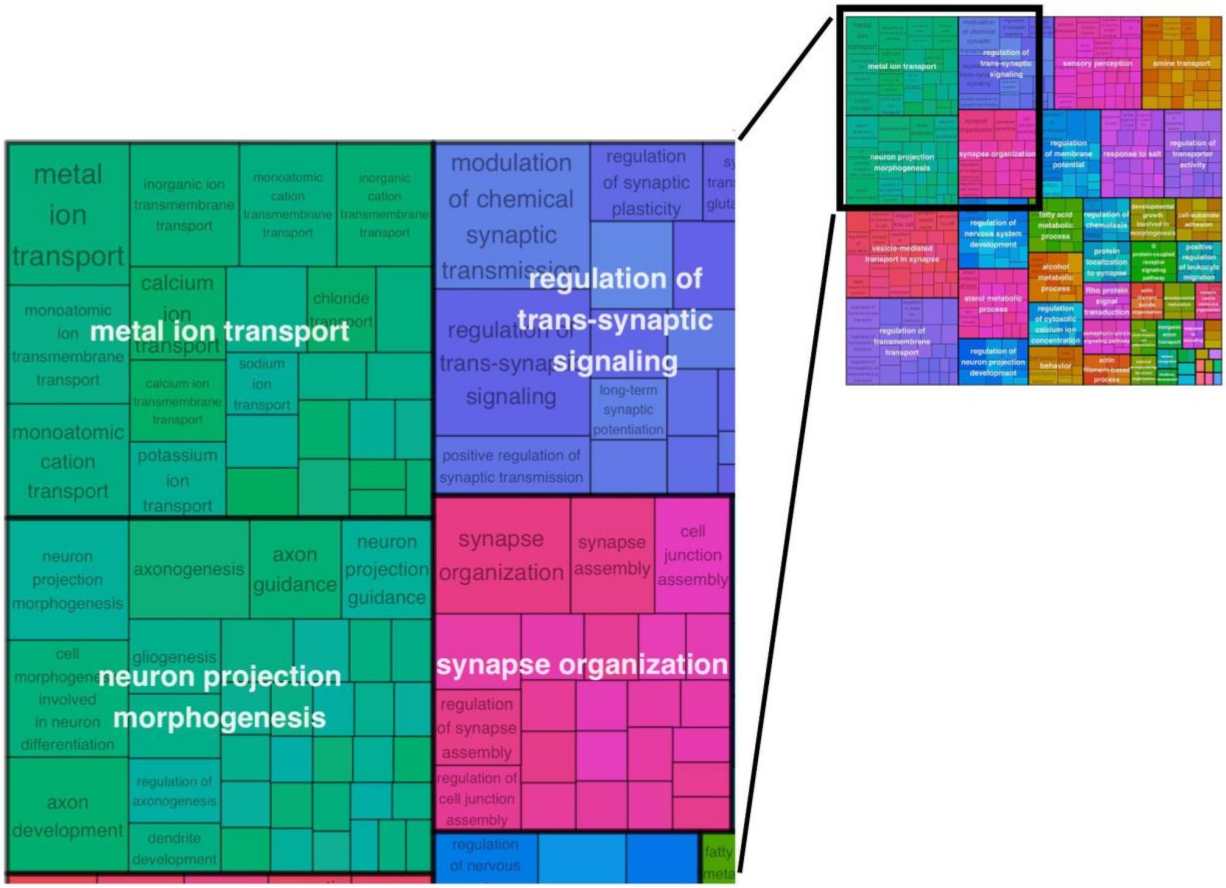


**Fig. S8. GO enrichment summarization by treemap.** Treemap of the enriched GO terms from the oligodendrocyte-versus-neuron comparison in brain tissue (Tabula Muris) — the same enrichment shown in Fig. S6 and summarized as a scatterplot in Fig. 1c and Fig. S7. Redundant terms are reduced by semantic similarity with rrvgo and grouped into clusters: each tile represents a GO term grouped under its representative (parent) term, tiles are coloured by parent term, and tile area is proportional to the term's significance (−log10 adjusted p-value). This space-filling layout places functionally related terms together, giving a compact overview of the dominant biological themes and complementing the scatterplot view.

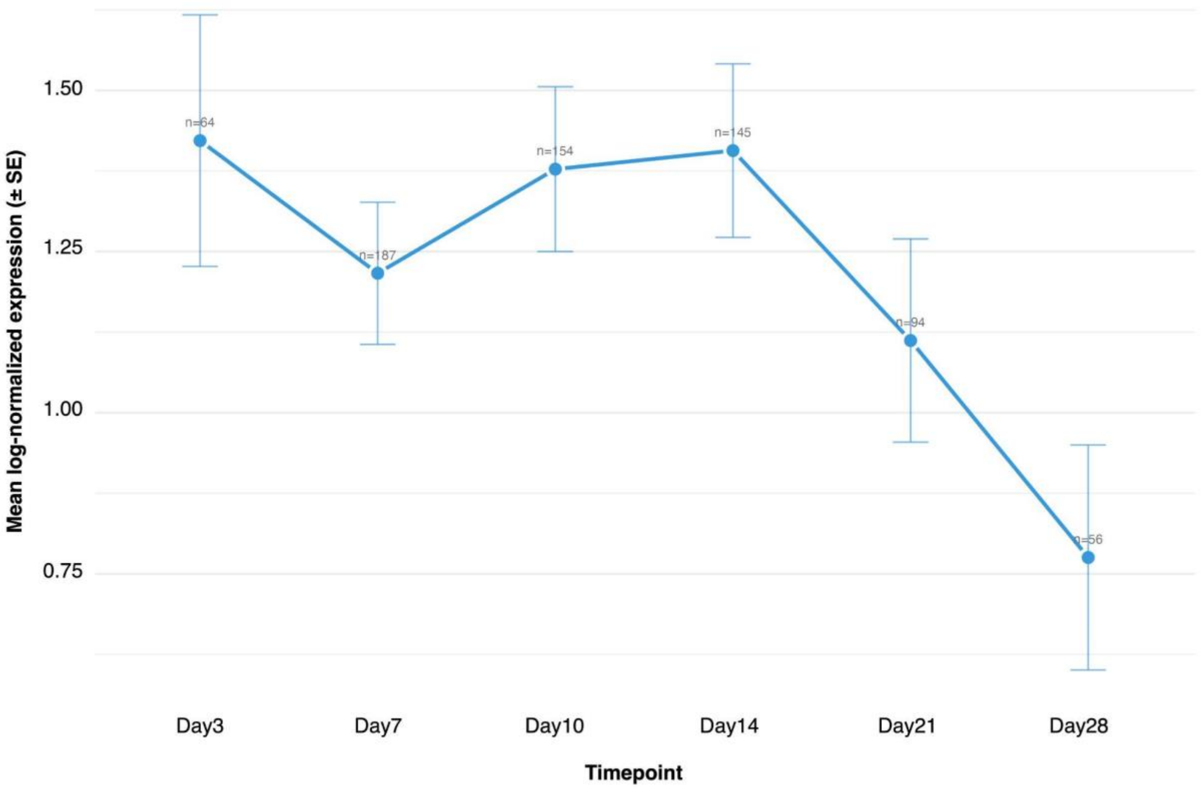


**Fig. S9. Temporal expression trend of Krt8 in Krt8+ ADI cells.** Mean log-normalised Krt8 expression in Krt8+ alveolar differentiation intermediate (ADI) cells across the bleomycin lung-injury time course (days 3, 7, 10, 14, 21, and 28). Each point is the per-timepoint mean across all profiled cells; error bars denote ± standard error, and *n* indicates the number of cells per timepoint. Krt8 expression remains high through the early-to-mid injury phase (days 3–14) and declines progressively thereafter (days 21–28), consistent with resolution of the transitional ADI state during later-stage repair. Generated with the Temporal Trend module of AtlasLens.

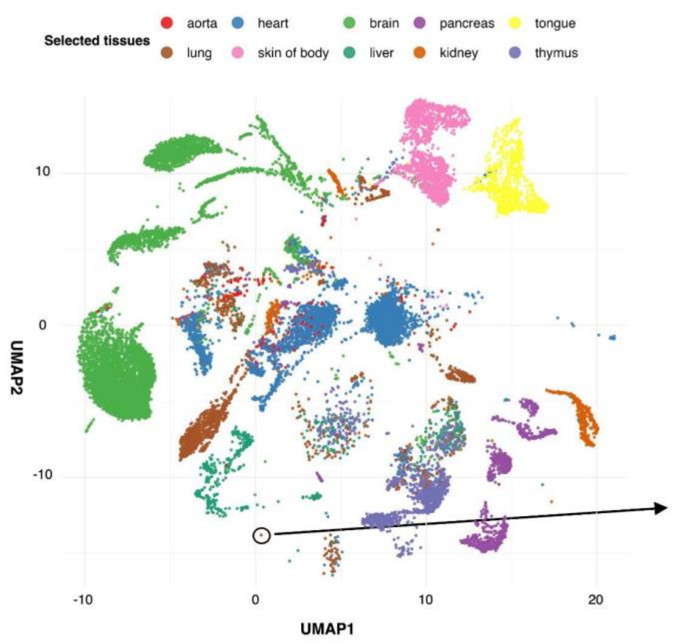


| Metadata field | Value |
|---|---|
| orig.ident | SeuratProject |
| nCount_RNA | 2942 |
| nFeature_RNA | 1360 |
| FACS.selection | Multiple |
| age | 18m |
| cell | P12_B002843 |
| free_annotation | Neutrophil |
| method | facs |
| donor_id | 18_47_F |
| subtissue | EPCAM |
| n_genes | 1429 |
| n_counts | 474900 |
| louvain | 13 |
| leiden | 12 |
| assay_ontology_term_id | EFO:0008931 |
| disease_ontology_term_id | PATO:0000461 |
| cell_type_ontology_term_id | CL:0000775 |
| tissue_ontology_term_id | UBERON:0002048 |
| tissue_original | Lung |
| development_stage_ontology_term_id | MmusDv:0000089 |
| self_reported_ethnicity_ontology_term | na |
| sex_ontology_term_id | PATO:0000383 |
| is_primary_data | TRUE |

**Fig. S10. Single-cell metadata inspection via interactive UMAP selection**. The cells shown are from the selected tissues in female 3 and 18-month-old mice from Tabula Muris dataset. A selected cell is highlighted on the UMAP, and all associated metadata for that cell are displayed in the table. AtlasLens supports interactive zoom and point selection to facilitate precise cell identification within dense embeddings.

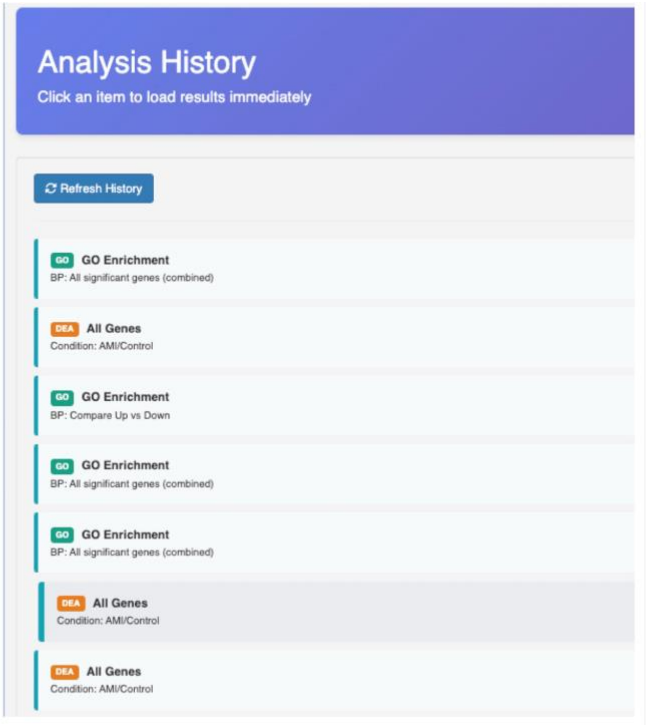


**Fig. S11. Session history panel in AtlasLens.** All analyses performed during an active session are logged and accessible in the History panel. Users can click any entry to revisit the corresponding analysis, inspect results, and download outputs, supporting reproducibility within and across sessions.
